# Transient states and barriers from molecular simulations and milestoning theory: kinetics in ligand-protein recognition and compound design

**DOI:** 10.1101/169607

**Authors:** Zhiye Tang, Si-Han Chen, Chia-en A. Chang

**Author notes:** The authors have contributed equally to this work (Zhiye Tang, Si-Han Chen).

## Abstract

This study applies a novel computational strategy to investigate molecular recognition and binding kinetics using five pyrazolourea ligands dissociating from cyclin-dependent kinase 8 with cyclin C (CDK8/CycC) as an example. The computed free energy barriers guide designing compounds using the transient conformations unavailable in experiments. The intermediates and their free energy profile during ligand association and discussion processes control ligand-protein binding kinetics and bring a more complete picture of ligand-protein binding. We used metadynamics and a pathway search method to sample pathways and applied combined reduced dimensionality, molecular dynamics (MD) simulations and milestoning theory to construct the free energy profile and estimate the residence time. The binding free energy and the trend of binding kinetics agreed with experiments. We explain the why of the barriers and the kinetics and use the information to assist ligand design. Guided by a barrier of a ligand passing an αC helix and activation loop, we introduced one hydroxyl group to parent compounds to design our ligands with increased residence time and validated our prediction by experiments. This work provides a novel and robust approach to investigate dissociation kinetics of large and flexible systems for understanding unbinding mechanisms and designing new small molecule drugs with desired binding kinetics.

**Significance Statement:** The transient conformations during ligand binding/unbinding control non-covalent binding kinetics. However, the transient structures and their free energy landscape of flexible ligand-protein systems are unavailable in experiments and challenging to model. Due to lack of understanding in binding kinetics, even scientists know that kinetic properties can be important in drug development, calculations using the intermediate states to design ligands with preferred binding kinetics are absent. We overcome these challenges and compute ligand-protein unbinding free energy profile using a novel method with molecular dynamics simulations, reduced dimensionality, and milestoning theory to deepen our understanding in molecular recognition. We also designed compounds based on the computed free energy barriers and experimentally validated that our designed compound can increase residence time.

## Introduction

Binding kinetics has become an important topic in molecular recognition because of the importance in fully understanding binding/unbinding and the growing awareness of the correlation between kinetics and drug efficacy^1-5^. Drug binding residence time, which can be estimated by a dissociation rate constant, 1/*k*_*off*_, is particularly important for determining the efficacy and selectivity of drug candidates. Experiments provide measured binding affinities (ΔG), rate constants (*k*_*on*_ and *k*_*off*_) and molecular structures. However, details are not fully presented by the experimental values and static conformations. As well, why a compound can bind/unbind fast or slowly is not fully understood. Molecular simulations, which are able to provide atomistic descriptions of temporal and spatial details of ligand–protein association and dissociation processes, become an important tool to characterize mechanistic features of binding kinetics and further assist drug development^6^. Features that govern binding kinetics are system-dependent and include ligand properties, conformational fluctuations, intermolecular interactions, and solvent effects^7-12^. However, the determinants to adjust for optimizing kinetic properties for a drug discovery project are not well understood.

All-atom molecular dynamics (MD) simulation in explicit solvent has been widely used to investigate protein dynamics and function as well as ligand–protein binding affinity. However, ligand binding/unbinding processes can be excessively longer than microsecond simulation lengths by using computer hardware available to most scientists^13-15^. Various methods such as aMD, metadynamics, weighted ensemble, scale-MD, and PaCS-MD have been used to accelerate sampling the ligand binding/unbinding processes^16-27^. In terms of the binding paths found, various algorithms such as umbrella sampling^28, 29^ and milestoning^30-33^ have been used to estimate kinetic rates and free energy profile. Multiple states may be identified from the trajectories, and the Markov state model (MSM) can be applied to estimate the transition rates^34, 35^. Using a reaction coordinate to accurately present ligand-unbinding free energy barriers provides invaluable information to understand binding kinetics and the mechanism. However, the task becomes daunting when a ligand–protein system under study is large and flexible. All important degrees of freedom involved during the unbinding processes must be included, but the use of high dimensionality to construct the ligand-unbinding free energy plot is impractical with umbrella sampling or milestoning theory. Therefore, this work developed strategies to obtain variables that cover important degrees of freedom for constructing a ligand-unbinding free energy profile with milestoning theory.

Cyclin-dependent kinase 8 (CDK8) is a promising cancer drug target because of its vital role in regulation^36^. CDK8 forms a complex with cyclin C (CycC), Med12, and Med13 for phosphorylation involved in positive and negative signaling of transcription and regulation of transcription activities. ^37, 38^ Abnormal activities of CDK8 and its partner CycC are implicated in various human cancers. ^39^ CDK8 has an allosteric binding site adjacent to the ATP binding controlled by a DMG motif (Asp-Met-Gly) that characterizes the DFG-in/DFG-out conformations as in other protein kinases. ^40^ A series of CDK8 drug candidates was discovered by structure-based drug design and virtual screening. ^41, 42^ A series of pyrazolourea ligands (PLs) have been developed and targeted the allosteric binding site of CDK8, ^43^ and a few studies have used MD simulations to examine the structural stability and ligand binding of the CDK8/CycC complex^44, 45^ A recent work used a metadynamics-based protocol to successfully rank the experimental residence time of CDK8 inhibitors^46^. The screening method provides a tool to identify inhibitors with relative short or long residence time; however, further investigation to understand determinants with atomistic details that govern the binding kinetics is needed.

In this work, we sampled dissociation pathways by using metadynamics and the newly developed pathway search guided by the internal motions (PSIM) method^47^. The free energy profile and residence time for five PLs shown in Figure 1 were computed by using milestoning theory with a novel algorithm to define the milestones. The work developed a new strategy involving principal component analysis (PCA), a mathematical method that can be used to extract the major motions from a collection of data, to define unbinding coordinates for constructing unbinding free energy barriers by using milestoning theory. By projecting the ligand unbinding trajectories onto the first two principal components (PCs), we were able to provide an unbinding coordinate for our milestones that considers 3N-6 degrees of freedom to capture the most important degrees of freedom involved during ligand unbinding, where N is the number of atoms. Different from MSM, which usually considers a handful of discrete states, we used more than 100 milestones to reveal smooth and detailed molecular motions and interactions corresponding to the unbinding free energy barriers. The computed free energy profile for ligand dissociation clearly indicates and explains where and why the energy barriers occur, such as important interaction formations/breakages between the ligand and CDK8, and motions of the ligand, CDK8 and CycC. Because the R-group of the PL compounds forms stable van der Waal contacts with CDK8, the R-group does not lead to ligand dissociation. Instead, the R-group serves as a hinge that allows the functional group to rotate and direct ligand unbinding. We found that the intermolecular hydrogen bonds (H-bonds) in crystal structures are critical to maintain the ligand binding mode in a bound state, and breaking a key H-bond cost ∼1 kcal/mol, which is in the range of H-bond strength computed from existing calculations. However, the major barriers arise from the concurrent motions of ligands and opening the protein binding site, which result in less favorable intermolecular attractions. We suggest the use of a bulky or hydroxyl group right next to the R-group of PL1 and PL4 instead of a linear alkane to increase their binding residence time. The calculations also provide the lower limit of the residence time, on a time scale of milliseconds and microseconds, and the trend agreed with experiments as well. Our calculations estimated that adding a small hydroxyl group increases by ∼2 to 3 times the binding residence time. The new PL4-OH compound was synthesized for experiments, and the kinetic assay validated our design and prediction. Guided by unbinding free barriers, the work introduces a new computer-aided design approach to modify compounds for preferred kinetic properties.

**Figure 1.**
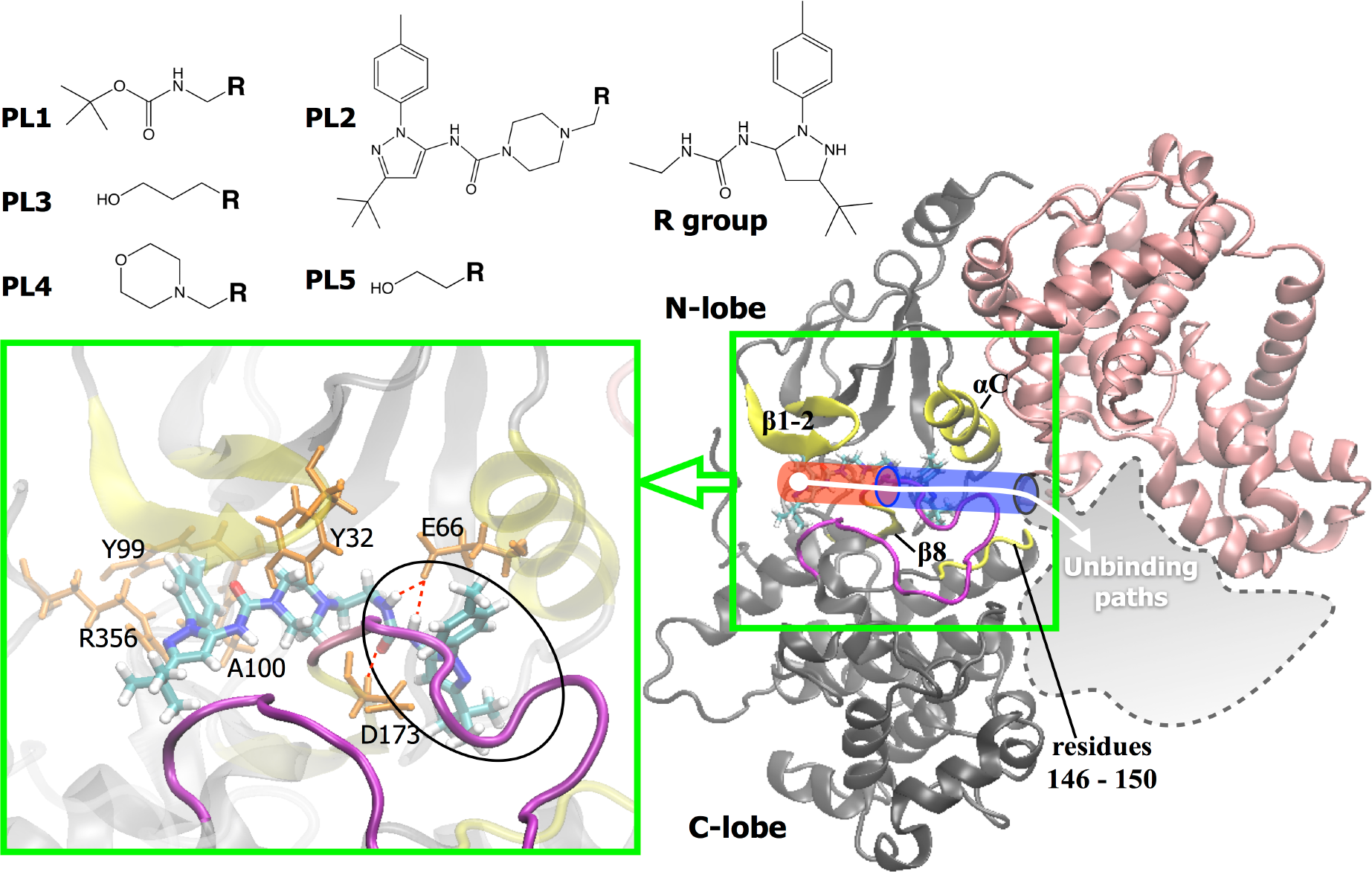
Protein structure and five PL compounds used in this study. Right: CDK8 (gray) and CycC (pink); left: a close-up view of ligand PL2 shown in licorice bound to CDK8. Residues engaged in important interactions with PLs in the bound complex are labeled with one-letter amino acid codes (orange). Conserved H-bonds between the urea linker of all five compounds with Glu66 and Asp173 are shown in red, and the circle indicates the R-group of PL compounds. Three stages of the dissociation process of the PL are represented as a red cylinder (leaving the front pocket, stage 1), a blue cylinder (passing a gate of the deep pocket, stage 2), and a gray amoeba (surface diffusion step, stage 3). Regions with significant motions during ligand unbinding are presented by different colors. Yellow: αC helix, β1, β2, and β8 sheets, and residues 146-148. Magenta: activation loop.

## Results and Discussion

### Uncovering energy barriers during dissociation using molecular simulations

Figure 2 illustrates complex free energy barriers along milestones that present important protein motions and ligand unbinding. Sampling ligand dissociation is the first critical step in understanding kinetic mechanisms of ligand dissociation. We used metadynamics, an enhanced MD simulation method, to obtain multiple ligand dissociation pathways for each compound. We also used the PSIM method to sample dissociation pathways. From our dissociation trajectories, we observed several kinetically relevant features, such as the first requirement for unbinding a ligand is breaking the key H-bonds that anchor the ligand to the specific binding modes and an αC helix and/or β1-2 sheet must move outwards. Then the milestoning theory was used to further quantify the free energy barriers associated with the conformational rearrangements of both the ligand and protein. Comparing the free energy profiles of the fast and slow ligands can lead to the design of new drugs with desired unbinding kinetics.

**Figure 2.**
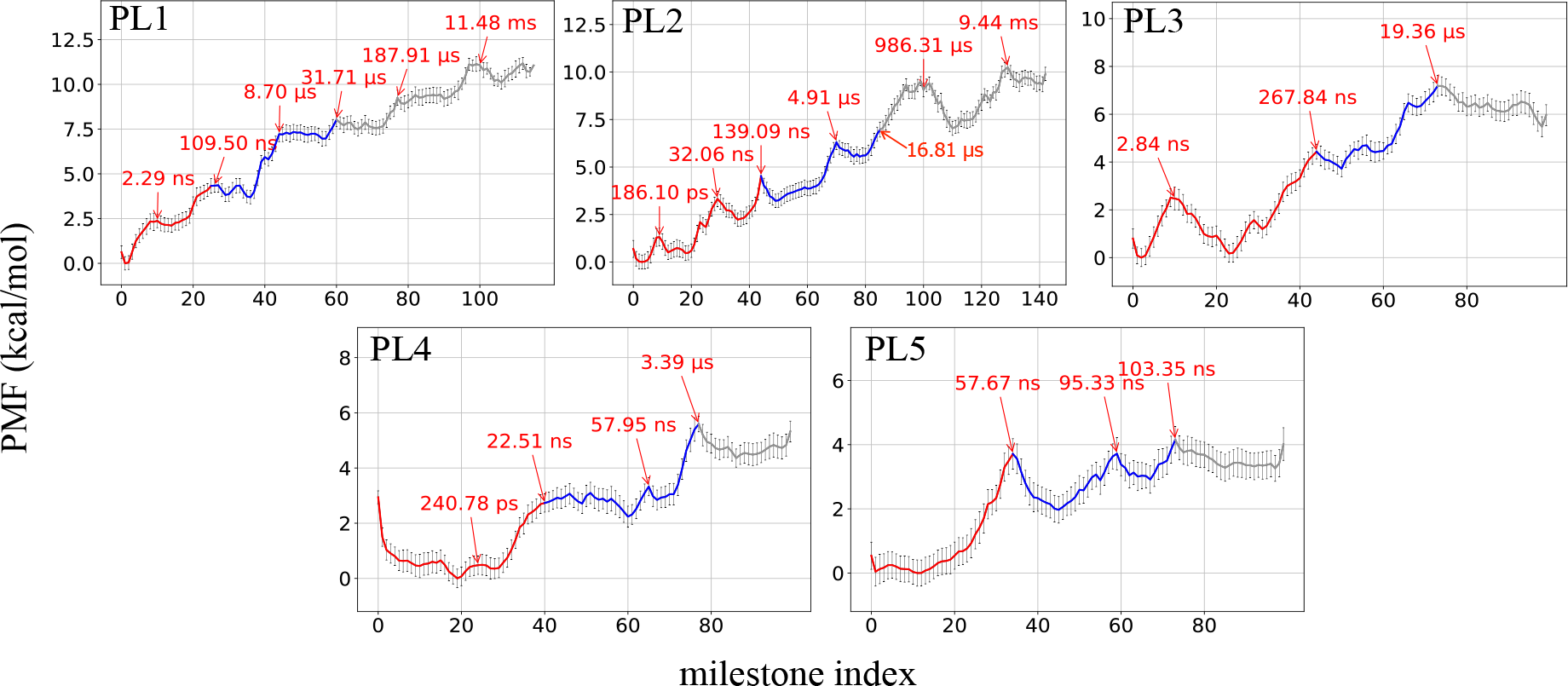
Free energy profiles along milestones during dissociation of CDK8/CycC-PL complexes. The transition times required to travel from the bound state to a barrier are marked in red. Initial barriers for breaking the conserved H-bonds between PL compounds and Glu66 or Asp173 and escape from the front pocket are colored in red (stage 1). All the PL compounds pass the gate of the deep pocket formed by Arg65, Trp146, Leu148, and Arg150, which resulted in energy barriers in blue (stage 2). In stage 3 (gray regions), PL compounds take diverse surface diffusion pathways as illustrated in Figure 1.

Because the dissociation process is complicated, manually selected states or milestones can create artificial bias. Therefore, we developed a novel strategy that uses natural molecular motions illustrated by PC modes from a dissociation trajectory to represent the unbinding coordinate of the milestones along a dissociation pathway (Movies S1 and S2). Here we used dissociation pathways obtained by metadynamics, and the strategy can be used for any trajectory with other modeling methods. The first two PC modes can represent more than 73% of protein motions (Figure S1). Because not 100% of protein motions can be presented by the first two PC modes, some small energy barriers may inevitably be missed in our dissociation free energy profile. However, because the PC modes capture major motions during ligand dissociation, all the important energy barriers should have been included in the free energy profile.

When a ligand is unbinding from the cavity, it can visit various regions on the protein surface before finally dissociating into the solvent. Because the surface diffusion step is not critical in determining a fast or slow binder, it is not a focus in the current study. For all CDK8/CycC-PL complexes we studied, the first stage of dissociation processes aims to break the conserved H-bonds between the ligand and the protein; as a result, the hydrophilic moiety can leave the front pocket (red cylinder in Figure 1, stage 1). At the next stage, the ligand keeps moving outward until it passes the gate of the deep pocket (blue cylinder in Figure 1, stage 2). Finally, diverse diffusion routes appear before the ligand completely escapes from the protein, which is partly captured in our pathways (gray amoeba in Figure 1, state 3). Therefore, the free energy difference between the milestones of bound states and the barriers to break the interactions with Arg65, Trp146, Leu148, and Arg150 are used to benchmark binding affinities (i.e., the difference between the minimum and maximum in the free energy profile, excluding the gray region). The last few milestones (Figure 2: gray regions) are part of the surface diffusion step, which is not included in this study because of incomplete sampling of the surface diffusion steps. Completely unbound states that include free translation and rotation of the ligand is not included in the free energy profile either. The computed relatively free energy, ΔΔG, agrees with experiment (Table S1). Because of the incomplete sampling for surface diffusion steps and ignoring the ligand free state with free translational/rotational diffusion, a larger absolute ΔG in our free energy profile than the experimental value is anticipated. The milestoning method revealed that ligands PL1 and PL2 have more rugged dissociation free energy profiles and more high-energy barriers to overcome during unbinding (Figure 2). The computed residence time and ΔΔG reflected the trends from experiments, and the computed residence time of PL1 and PL2 was an order of magnitude slower than the ligands, PL4 and PL5, which clearly reveals the significance of barriers in stages 1 and 2 during the dissociation process. In addition, the average time required to cross each notable barrier can be estimated (labeled in Figure 2).

### Identifying important dissociation steps and the structure–kinetic relationship

The energy barriers illustrated by our free energy profiles reveal various protein conformational rearrangements, loss of intermolecular attractions, and changes of the H-bond network during ligand unbinding. For example, intermolecular H-bonds between the urea linker with Glu66 and Asp173 must be broken for unbinding this series of PLs (Figure 1); however, this step does not yield the same free energy cost for every ligand. PLs jiggle in the bound state, and local molecular fluctuations can temporarily break an H-bond, which costs ∼1 kcal/mol in our free energy plot. Only permanently breaking H-bonds between the urea linker with Glu66 and Asp173 permits PL compound dissociation, which requires significant molecular rearrangements. For example, in the snapshot of barriers A and B in Figure 3, positions of αC, β1-2, and β8 are adjusted in order to pave a pathway for PL1 to exit.

**Figure 3.**
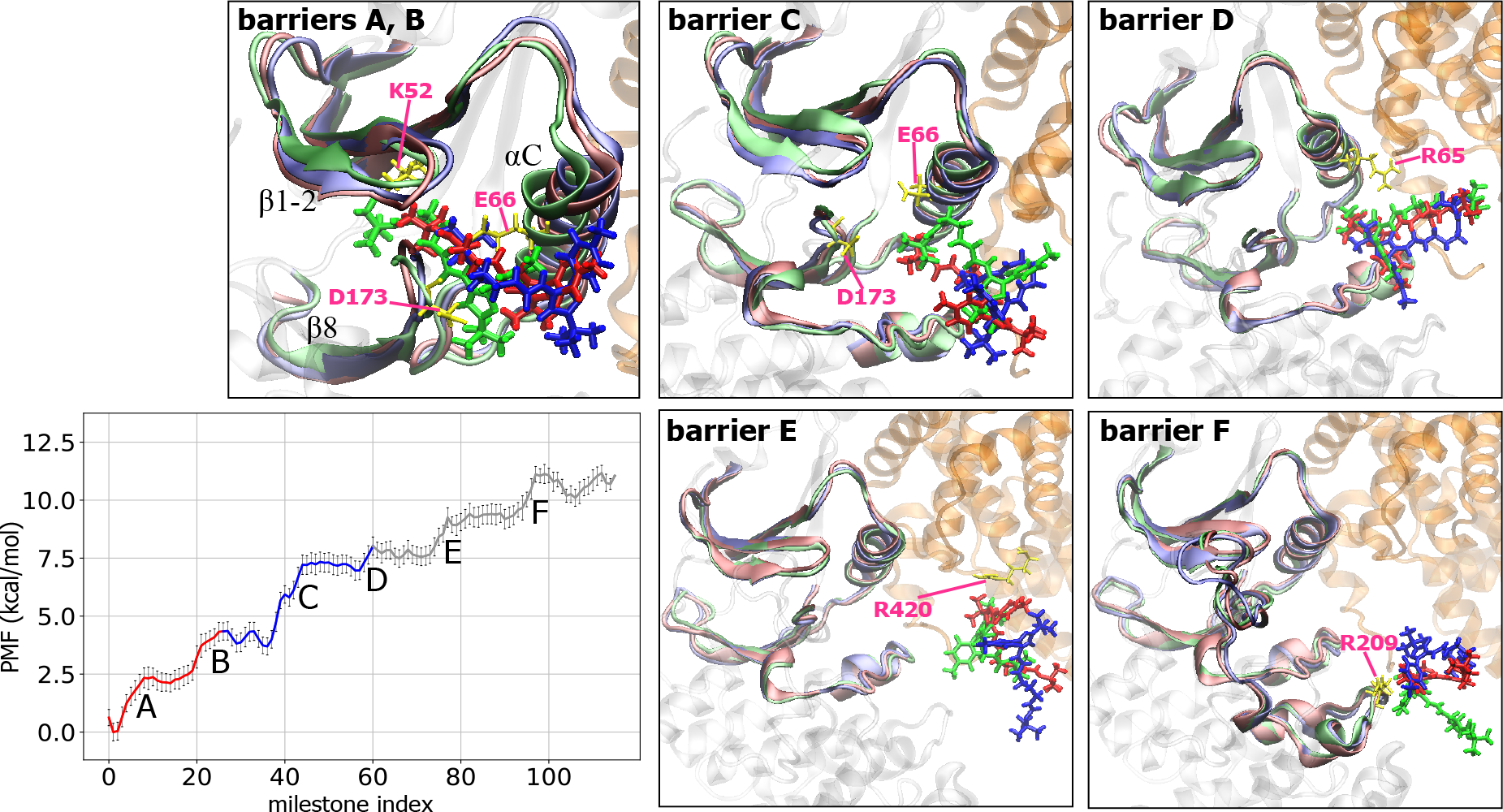
Free energy profiles along milestones and important conformational changes associated with each major energy barrier during PL1 dissociation. See Figure 2 legend for the details of the free energy profile. Snapshots for each labeled energy barrier illustrate conformational changes of CDK8/CycC-PL1 when PL1 passes. PL1 is shown in licorice. CDK8 and CycC are colored in gray and orange, respectively. For each figure, initial, halfway and final conformations are shown in green, red, and blue, respectively, for both PL1 and the binding pocket, and key residues in the initial conformation are labeled and represented as yellow sticks.

Although the most important stage in molecular dissociation is the initial step, during which a compound mostly breaks crucial intermolecular interactions in the bound state and starts to leave the binding pocket, it is not the whole picture of binding kinetics. The intermediates affected by the structure and properties of a compound may largely contribute to the free energy barriers and binding kinetics as well. Notably, the energy barrier associated with this important initial stage is not related to binding free energy either (Figure 2). PL1 has a second set of polar linkers, which forms additional H-bonds with Lys52, Glu66, and Asp173 at the bound state and increases the energy barrier height and transition time to overcome the barriers A and B in Figure 3. The H-bond donor and acceptor of PL1 also display multiple formation and breakage of complex H-bond networks with CDK8 and/or the bridge water molecules in subsequent milestones, as illustrated in Figure S6. The interactions contribute to stable intermediate states (snapshots in Figure 3) and multiple energy barriers to leave these local energy minima. In contrast, the morpholine group of PL4 is not able to form strong attractions with CDK8, which results in less rugged free energy barriers during PL4 unbinding (Figure 2).

PL2, the largest of the five ligands, shows similar behavior to PL1: stepwise H-bond formation and breakage, significant molecular rearrangements in stage 1, and multiple subsequent energy barriers during unbinding (Figure 4, and S2). Molecular fluctuations may temporarily break the H-bonds between Glu66 and Asp173, which result in two small energy barriers (∼1 kcal/mol) before heading to a major energy barrier (Figure 4: barrier B). Notably, although unbinding free energy profiles computed from different dissociation pathways are not identical, the key events are the same. For example, the first major peaks in both Figures 4 and S2 denote the free energy cost for permanently breaking the H-bonds between Glu66 and Asp173, and the energy cost of escaping the front pocket (stage 1) from both pathways is ∼4 kcal/mol. However, PL2 can have slightly different fluctuations before breaking this major interaction, which results in minor variation in the free energy plots.

**Figure 4.**
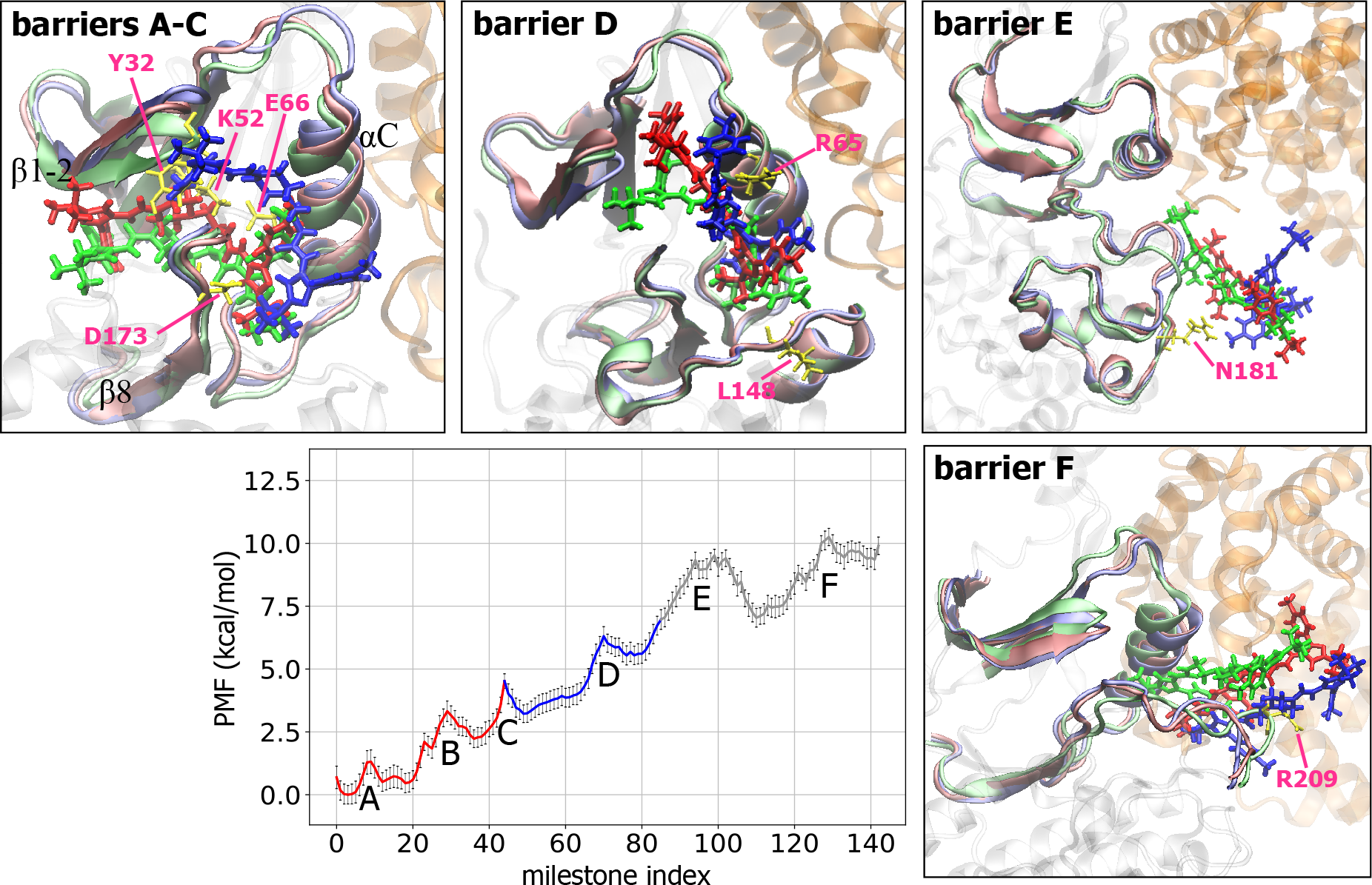
Free energy profiles along milestones and important conformational changes associated with each major energy barrier during PL2 dissociation. See legends in Figures 2 and 3 for details.

PL3 and PL5 are the two smallest compounds, with a linear 5-hydroxypentyl and 4-hydroxybutyl group, respectively. Besides the conserved interactions, at the bound state, PL3 is stabilized by the H-bond between the 5-hydroxypentyl group and Asp98 in the hinge of CDK8. Similar to PL1 and PL2, both protein rearrangements and step-wise breakage of the H-bonds resulted in the rugged free energy profile at stage 1 (Figure S3). In contrast, the 4-hydroxybutyl group of PL5 could not reach the hinge region, and the smaller PL5 escaped from the binding pocket without invoking large protein rearrangements (Figure S4: barrier A). Interestingly, after passing the initial unbinding barriers, all PLs retained strong interactions with the αC helix by forming new H-bonds with Arg65. While the compounds are unbinding, the free energy continues to increase until the end of stage 2, when the interaction with the αC helix vanishes in all free energy profiles. Once exiting the gate of the deep pocket, the compound starts to diffuse over the surface of the CDK/CycC complex, and several intermediates form before complete dissociation. For example, at barriers E and F in Figure 2, PL1 interacts with CycC and the C-lobe region of CDK8, respectively, and at barrier E in Figure 3, PL2 forms a stable H-bond with the highly fluctuating activation loop of CDK8. Of note, a handful of pathways for the diffusion steps are not sufficient to fully describe ligand surface diffusion, which needs a statistical ensemble of ligand diffusion trajectories. Importantly, milestones of stages 1 and 2 inform both the structure–kinetic and structure–activity relationships (Table S1).

### Understanding unbinding mechanisms of fast (PL4) and slow (PL2) dissociation

Here we investigated the unbinding paths and their intermediates in greater detail for one slow (PL2) and one relatively fast (PL4) binding/unbinding compound. We selected these two compounds because of the significant differences in their hydrophilic moieties, with PL2 forming additional H-bonds through the second set of urea linker and pyrazolyl nitrogen atoms. In contrast, PL4 has no good H-bond donors or acceptors besides the R group.

PL2 is the largest compound in this study and is anchored by the H-bonds between Glu66 and Asp173; the other functional groups all form nice contacts with the kinase front and deep pocket (Figures 1 and 4). At energy barrier A, the hydrophilic moiety of PL2 begins to jiggle inside the front pocket, due to the weakening of several interactions between PL2 and the binding pocket, for example, water-bridged H-bonds with Val27 in β1-2, Ala155 and Asn156 in the front pocket and Arg356 in close proximity of the hinge. We also observed the same molecular motions and disruption of interactions from other unbinding pathways for PL2 (Figure S2), although not all local energy minima are identical during unbinding. The major time-limiting step during stage 1 is the energy barrier B, where the stable H-bonds between PL2 and residues Tyr32, Glu66, and Asp173 permanently break. Together with the upward movement of β1 and β2 sheets, the 3-tert-butyl-1-(4-methylphenyl) group rotates, which allows the ligand to leave the binding site (Figure 4: barriers A-B). During this step, the piperazine ring located in the center of PL2 passes the cleft formed by β1/β2 sheets, β8 and the activation loop, which requires protein arrangement for opening the cleft. At energy barrier C, the β1/β2 sheets keep moving upward until the hydrophilic moiety of the PL2 completely leaves the cleft, together with breakage of the H-bonds between PL2 and the β1/β2 sheets. As compared with PL4, during PL2 unbinding, the upward movement of the β1/β2 sheets is more pronounced because the β1/β2 sheets can form more contacts with this large ligand and result in concurrent motion with PL2 (Figure 5). Instead of having multiple small barriers during the initial stage, PL2 can also break the conserved H-bonds with Glu66, Asp173, the β1/β2 sheets within one major energy barrier (e.g., barrier A in Figure S2). Both dissociation pathways require ∼4.0 kcal/mol for PL2 to completely pass the cleft (Figure 4: barrier C and Figure S2: barrier A), with transition times 139.1 and 101.1 ns, respectively. Because of only one possible channel for unbinding the PL compounds, the unbinding kinetics of stage 1 could be highly similar, which is supported by our examination using independent runs of PL2.

**Figure 5.**
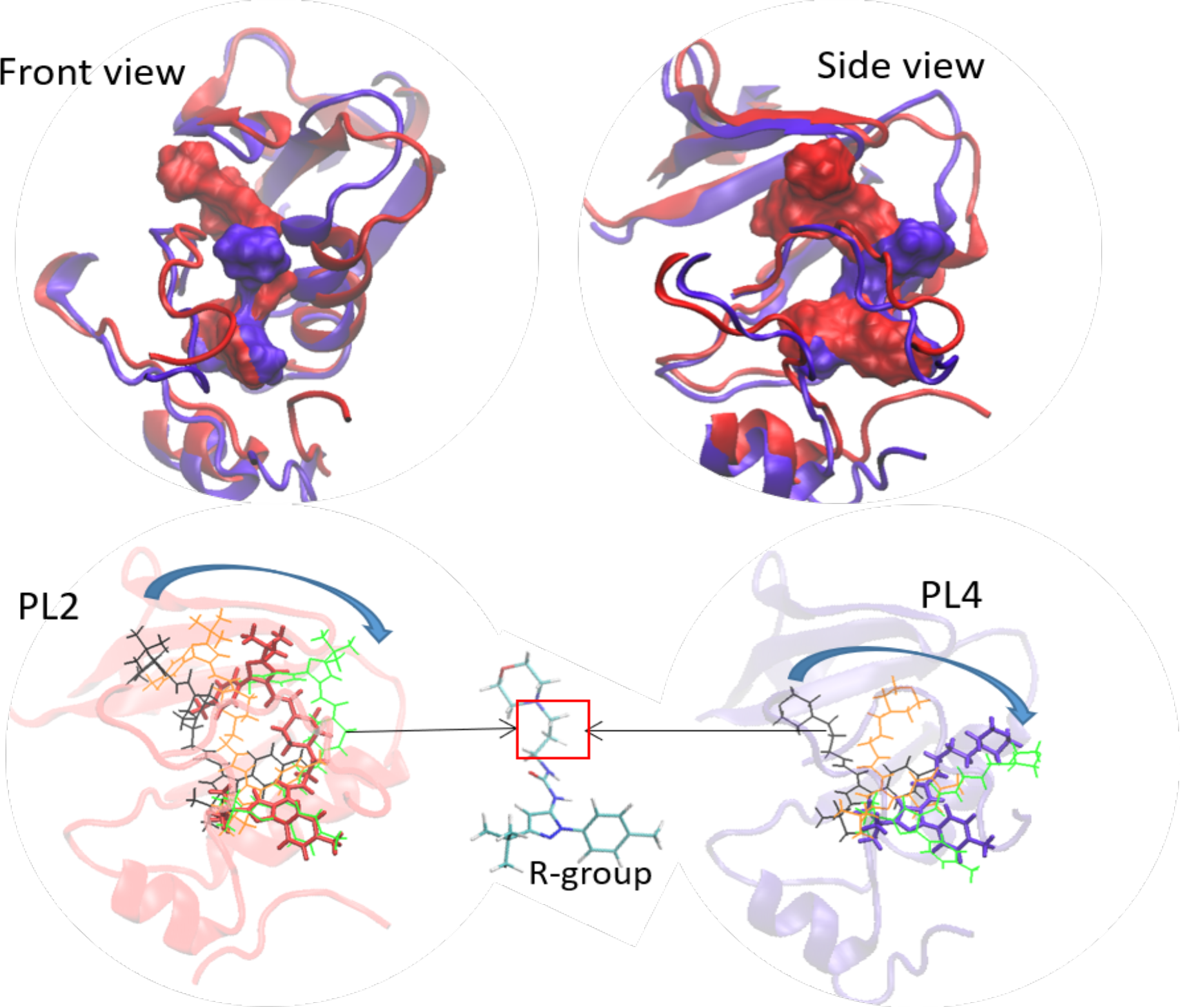
Conformations of PL2 (red) and PL4 (purple) during dissociation. Top: conformations when PL2 and PL4 are passing barrier B in Fig 4 and B in Fig 5, respectively. Nobly, PL2 is significantly larger than PL4, so unbinding PL2 requires more open β1 and β2 sheets and αC helix. Bottom: hinged by the R-group, the ligands rotate the functional groups during dissociation. PL2 moves from a stable bound state (black) to barrier B (orange), passing barrier B (red) and barrier C (green). PL4 moves from a stable bound state (black), to passing barrier A (orange), middle between barriers A/B (purple) and barrier B. PL4 is presented in the middle, and the square indicates an alkane chain that may be modified by a bulky and/or polar group to increase the residence time of PL4.

After overcoming the unfavorable step, a new H-bond network between the compound, protein and bridge water molecules forms, which leads to an intermediate local energy minimum after energy barrier C in Figure 4. When the urea linker of PL2 anchors the αC helix through the H-bond with Arg65, the pyrazolyl nitrogen of the R-group also forms a water-bridged H-bond with Leu148 at the linkage between αE and αF helices, thereby generating a stable intermediate. Breaking the interactions with Leu148 and αC helix leads to energy barrier D in Figure 4 and barriers B-D in Figure S2. As shown in Figure S7, the network of the H-bond network between PL2 and Arg65 survives up to the left shoulder of barrier E in Figure 4. Therefore, in both PL2 dissociation pathways, the barriers in stage 2 take ∼20 to 30 µs and are critical steps for other compound dissociations. The major motions all appeared in our metadynamics and PSIM runs (e.g., Figures 4, S2 and S10). After leaving the ligand binding site, PL2 continues forming and losing interactions with CDK8 and CycC (Figure 4E/F) until it is fully solvated by water molecules. The simulations also show the breathing motion between the N- and C-lobes of CDK8, whereby the two lobes move apart to open the binding sites for ligand dissociation and then shift to come closer after PL2 completely leaves CDK8 (Figure 4, Milestone Index 140). We observed the same breathing motion during classical MD simulations for apo CDK8/CycC proteins^45^. The breathing motion was also found closely related to the ligand binding/unbinding from p38 mitogen-activated protein kinase^7^.

The computed free energy profiles explain why inhibitors such as PL4 demonstrate fast unbinding kinetics by revealing the heights and numbers of free energy barriers. As for all other inhibitors in this series, the bound state of PL4 is stabilized by the two H-bonds via the urea linker of type-II ligands with Glu66 on the αC helix and Asp173 on β8. However, the terminal [3-(morpholine-4-yl)propyl] group forms relatively weak intermolecular interactions with the hinge residues, Ala100 and Asp98. At stage 1, the interactions between PL4 and the hinge loosen and the [3-(morpholine-4-yl)propyl] group escapes from the front pocket after breaking the H-bonds between the urea linker and residues Glu66 and Asp173. However, PL4 cannot leave the pocket unless it pushes the β1/β2 sheets to move upward, thus creating room for further dissociation (Figure 6: barrier A, and Figure 5). This step features a < 1 kcal/mol energy barrier, which suggests that the rearrangement of the binding site is easy. The barrier-less flipping of the morpholine group is mainly attributed to the relatively small hydrophilic moiety of PL4, making a more flexible binding pocket. This finding is consistent with the classical MD simulations of CDK8/CycC-PLs at the bound states in Figure S10, in which the CDK8/CycC-PL4 complex, as compared with other PLs, shows greater breathing motion between the C-lobe and N-lobe of CDK8. The intermediate after barrier A is stabilized by water-bridged H-bonds with Glu66 on the αC helix, Asp173 on β8, and Arg178 on the activation loop (Figure S8). The major energy barrier functions to permanently break this H-bond network (Figure 5: barrier B). Of note, the activation loop does not directly involve ligand binding affinity^45^, but the loop can affect binding kinetics.

**Figure 6.**
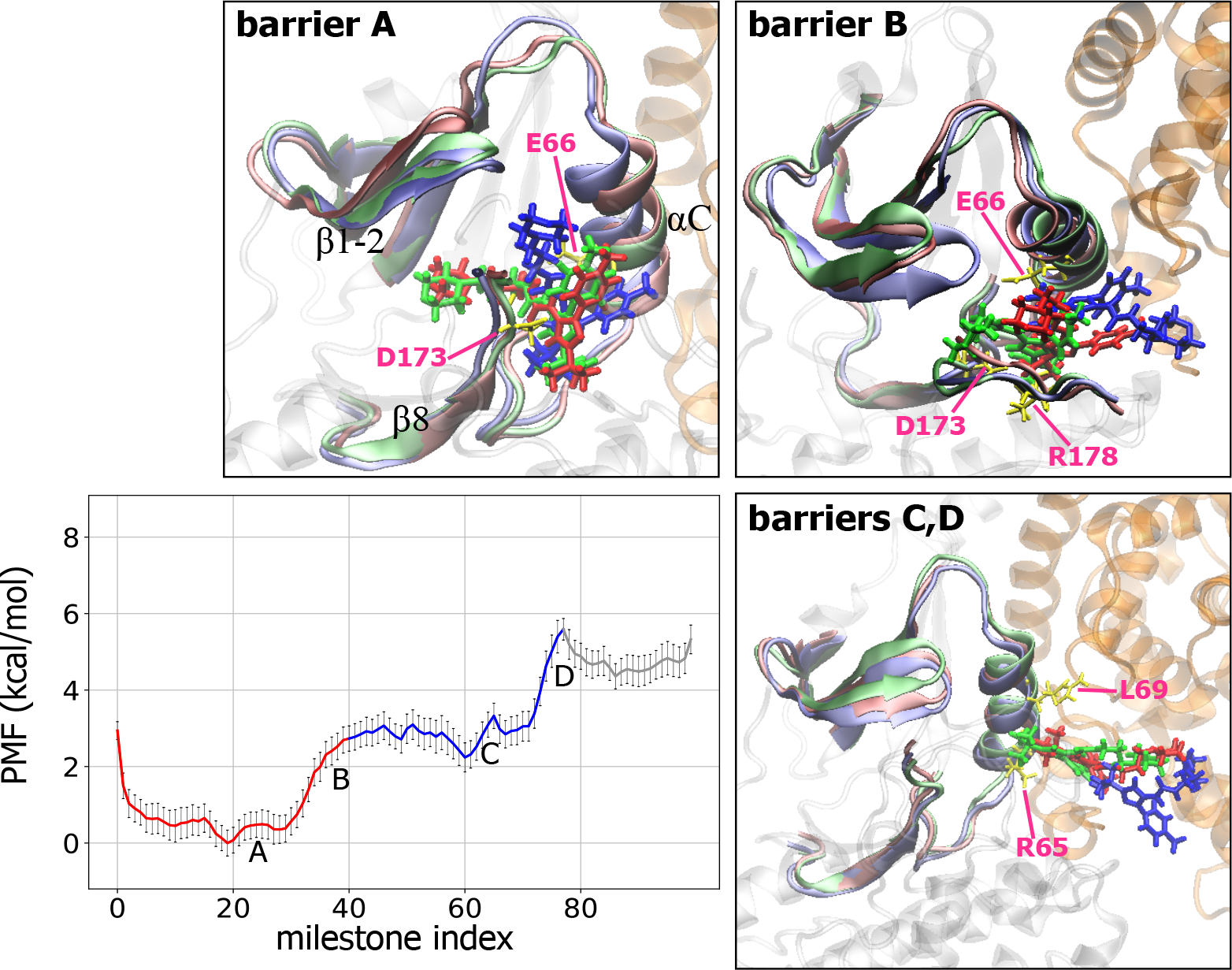
Free energy profiles along milestones and important conformational changes associated with each major energy barrier during PL4 dissociation. See legends in Figures 2 and 3 for details.

After stage 1, the energy keeps increasing before completely breaking the H-bond network between PL4 and both the activation loop and αC helix, thereby resulting in the last two major energy barriers C and D in Figure 6. Unlike the other PL compounds, PL4 cannot form a H-bond network with Trp146, Leu148, or Arg150 on passing the gate of the deep pocket, which results in a relatively short transition time for stage 2. Although the major unbinding events are similar, the lower free energy barrier heights of PL4 result in a faster estimated residence time than in PL2.

### Modifying PL ligands to increase binding residence time

The energy barriers associated with particular movements and/or interactions during dissociation inform rational drug design for desired kinetic properties. As illustrated in Figures 4 and S2, passing the second pyrazol and benzene rings of PL2 produces a high energy barrier as compared with the smaller [3-(morpholine-4-yl)propyl] group of PL4, which requires less significant motion of β1 and β2 sheets. Large ligands such as PL2 may demand more unfavorable protein motions to unbind the ligand, and our calculations show that non-specific van der Waal attractions can increase the cost of ligand unbinding (data not shown). However, a large ligand is not always desirable. Here we suggest introducing a bulky group such as naphthalene, benzopyran or benzofuran next to the R group of PL4 (Figure 5) to strengthen the non-specific attraction between the αC helix and activation loop when the ligand is passing the cleft. Another promising drug design is inspired by the results of PL3 and PL5, in which the hydroxyl groups can stabilize the bound state by forming additional H-bonds or holding a bridge water molecule with front pocket residues (e.g., Lys52, Asp98, and Ala100). More importantly, we found favorable interactions between the hydroxyl groups and residues 146-50 while the compounds pass the gate of the deep pocket, which increase the height or number of the free energy barriers after the initial step of dissociation. Therefore, adding a hydroxy in the alkane chain of PL4 or a carbonyl and/or hydroxyl group in the naphthalene ring next to the R group may further slow the unbinding process.

Following the above strategies, we designed ligand PL4-OH by introducing an additional hydroxyl group to the [3-(morpholine-4-yl)propyl] group of PL4 (Figure 7) and performed proof-of-concept calculations and experiments. The hydroxyl group of PL4-OH formed an H-bond with a front pocket residue, Lys52, at the bound state (Figures 7 and S9). However, this additional interaction had negligible contribution to the free energy barriers and transition time for stage 1 (Figure 6). After passing barrier A, the complex is stabilized by the newly formed H-bond between the hydroxyl group of PL4-OH and Asp173, which requires additional energy to break (Figure 7: barrier B). Upon passing the gate of the deep pocket, the two major barriers (Figure 7: barriers C and D) are attributed to step-wise breakage of the H-bond network between the hydroxyl group PL4-OH and Trp146, Leu148, and Arg150. As compared with other compounds, PL3 and PL5, the hydroxy-induced interactions between PL4-OH and CDK8 have more pronounced contribution to increasing unbinding residence time. This contribution could benefit from the relatively rigid [3-(morpholine-4-yl)propyl] of PL4-OH as compared with the linear hydrophilic moiety of PL3 and PL5. The computed residence time is 2.7 times longer than that for PL4. PL4-OH was synthesized and the experimental assays validated that the minor modification by adding a hydroxyl group successfully increased the residence time (Table S1).

**Figure 7.**
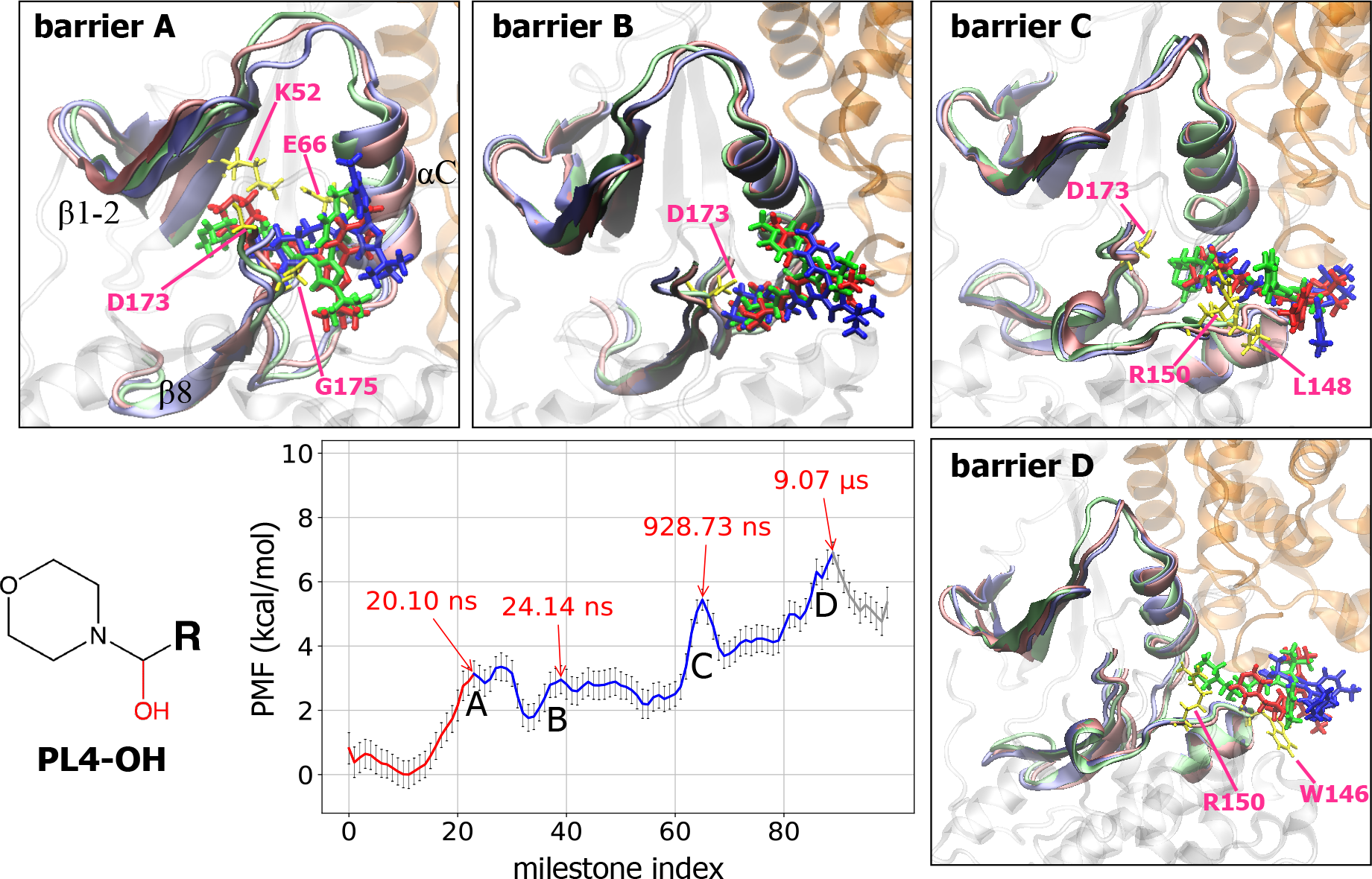
Free energy profiles along milestones and important conformational changes associated with each major energy barrier during PL4-OH dissociation. See legends in Figures 2 and 3 for details. The PL4-OH compound is shown as sticks with the additional hydroxyl group colored in red.

We further investigated this structure–kinetics relationship by performing calculations for the second designed compound, PL1-OH, with the same hydroxyl group added to PL1 (Figure S5). As expected, the hydroxyl group of PL1-OH forms an additional H-bond with Lys52 at bound state, which breaks together with the conserved H-bond with Asp 173 (Figure S5: barrier A). Before leaving the front pocket, the hydroxyl group forms a new H-bond with Asp173 and helps keep the compound inside the pocket. After breaking the conserved H-bond between the urea linker and Glu66 (Figure S5: barrier B), PL1-OH is stabilized by the H-bond network between the hydroxyl group and Leu148, which results in barrier C at the end of stage 2. Although the additional hydroxyl group participates in the H-bond network at the bound state, the longer residence of both PL4-OH and PL1-OH is mainly due to the increased number or stability of intermediates in stage 2. The studies demonstrate that using solely bound states is not sufficient to understand ligand binding residence time and guide ligand design with preferred kinetic properties, and a free energy profile along the dissociation pathway provides information on transient conformations for design not seen in experiments.

### Novel strategy, remaining challenges and future perspectives in free energy barrier-guided ligand design

Constructing an unbinding free energy profile needs a coordinate to present the unbinding pathway. Using a distance coordinate between a ligand and protein can miss the free energy contribution from rearranging protein side-chains and/or backbone. One may add a couple of dihedral rotations in the coordinate; however, key dihedrals change during ligand unbinding, and explicitly selecting certain degrees of freedom inevitably ignores motions that contribute to the free energy barriers. Therefore, we used molecular motions presented by the first two PCA modes, PC1 and PC2, to present the unbinding coordinate. The first two modes covered more than 73% of the major motions for all five ligands in this study (Figure S1). By using this novel approach to choose the unbinding coordinate for milestoning theory, we successfully obtained the free energy barriers that are useful for ligand design.

Figures 8 and S1 illustrate that using eigenvectors of PC1 and PC2 effectively reduces the dimensionality for presenting the unbinding pathway, not limited to a few collective variables such as the distance between two centers of mass or selected dihedral angles. Each dot denotes a frame from the unbinding trajectory. The red curve is the smoothed path, and we divided the space along the path into compartments separated by milestones (black line in Figure 8B). Milestoning theory, which follows statistical mechanics theory to estimate the transitions between compartments along the unbinding pathways, allows for using short trajectories initiated within each compartment. Notably, because of complex protein systems, initial structures for short MD runs are close to but not always exactly on the milestone interface. The trajectories are analyzed to produce free energy profiles and accurately estimate the time a ligand needs to pass each energy barrier and unbind.

**Figure 8.**
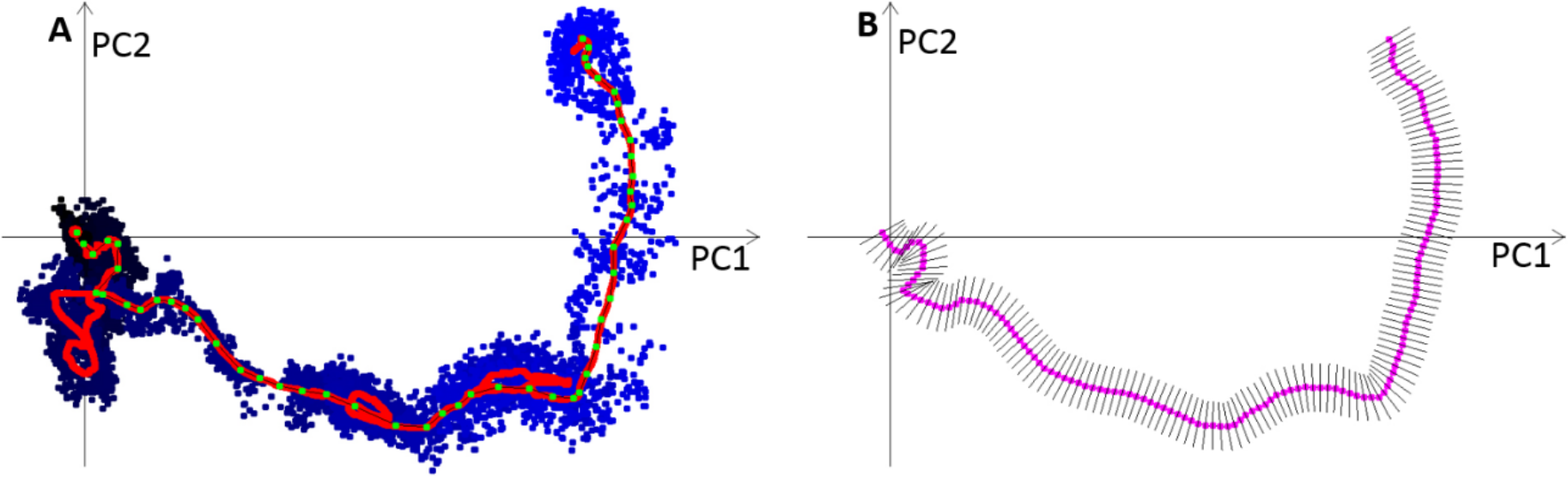
The projection of frames from a metadynamics trajectory of PL2 onto PC1/PC2 coordinates and spatial definition of milestones. A: the dissociation pathway projected on PC1/PC2 space (dots) and smoothed trajectory (red). Black and blue dots present the ligand position when it was inside or outside the protein binding pocket, respectively. The averaged position is illustrated in the red line as a smoothed trajectory. Green dots present the manually defined path based on the smoothed dissociation trajectory. B: the smoothed trajectory in A was optimized to remove the frames that do not lead to PL2 dissociation. The optimized path is in purple and the milestones are shown as black lines.

Because proteins have complicated molecular rearrangements, not all natural motions can be included in the compartments. Therefore, some fluctuations that contribute to unbinding free energy barriers are ignored, which results in faster kinetics. Introducing nonlinear dimensionality reduction algorithms in machine learning such as auto-encoders may further capture ignored small free-energy barriers that also contribute to the unbinding free energy profile. The transition matrix constructed by multiple short unbiased MD trajectories provides the transition probability between nearby milestones; however, the frequency of a ligand moving back and forth between numerous major or tiny energy barriers may not be highly accurately captured. The computed residence time may be considered the time required to smoothly travel from the bound to unbound state without excess backward and forward swing.

For tight binders with only one dissociation direction such as the system studied here, our calculations demonstrate that the initial movement during the unbinding process is highly similar from multiple pathways, and these initial steps determine the binding residence time and relative binding free energy. However, for protein systems with a wide-open binding site, such as HIV-1 protease, instead of one well-defined binding/unbinding channel, molecular modeling may sample significantly different dissociation pathways; thus, different unbinding directions need to be considered when constructing the unbinding free energy profiles. However, estimating the population of each pathway remains challenging. The milestones obtained from each pathway can differ, which increases the difficulty in combining multiple free energy profiles and selecting critical free energy barrier(s) for drug design. Additional theoretical work will provide more rigorous methods to assemble binding/unbinding free energy profiles from multiple pathways or determine important barriers for drug development.

## Conclusions

In this work, we performed a detailed investigation of pathways, binding kinetics, and thermodynamics for type II ligands unbinding from CDK8/CycC and used the free energy barrier to guide ligand design. We also performed experiments to validate our new design, PL4-OH, with increased binding residence time. This was achieved by first sampling dissociation pathways of a series of pyrazolourea ligands with diverse structures of the hydrophilic moiety, mapping high-dimensional protein–ligand dissociation pathways onto the reduced coordinates by using the first two PC modes to define milestones and applying milestoning theory to construct an unbinding free energy profile and estimate residence time. Ligand design is then guided by revealed rate-determining events, for example, substantial molecular rearrangements and breaking a conserved H-bond network for escaping the hydrophilic moiety of the front pocket of CDK8 (red line in Figures 2-4, 6-7) and multiple formation and breakage of new H-bond networks between a ligand and the gate of the deep pocket (blue line in Figures 2-4, 6-7). The study showed that the surface diffusion step (stage 3 in Figure 1) is not an essential factor to determine related residence times and ΔG, and focusing on the steps (stages 1 and 2) when a ligand is leaving the front and deep pocket can yield good agreement with experiments.

Our computed free energy barriers suggested that modifying the hydrophilic moieties of PL1 and PL4 would increase residence time. The calculations predicted 2 to 3 times increased residence time for adding one hydroxyl group to the parent compounds, PL1-OH and PL4-OH, which was validated by experiments. The longer residence time originated from the formation and breakage of H-bonds between the hydroxyl groups and residues at the gate of the deep pocket (Trp146, Leu148, and Arg150), which increases the number of intermediates and free energy barriers of the dissociation pathway. Successfully combining PCA and milestoning theory revealed detailed protein–ligand interactions/motions and unbinding mechanisms during the dissociation process and provided valuable structure–kinetics relationships for designing drugs with preferred binding kinetics.

## Methods

### Molecular simulations

We obtained the initial structures of CDK8/CycC–PL complexes from the PDB database (PDB ID: 4F6W, 4F7L, 4F6U, 4F7N ^43^) and manually modified PDB structure of 4F7N, 4F7L, and 4F6U to obtain the initial structures of CDK8/CycC-PL5, CDK8/CycC-PL1-OH, and CDK8/CycC-PL4-OH, respectively. VCharge^48^ was used to estimate partial charges of all ligand atoms. We added missing residues and built the missing activation loop of the structures by using Swiss Model ^49^ based on the p38 DFG-out crystal structure (PDB ID: 1W82 ^50^) as a template. We determined the protonation states of histidine residues in the CDK8/CycC–ligand complexes by using the MCCE package ^51^. AMBER FF14SB force field and GAFF ^52^ were used for the CDK8/CycC complex and PLs. We solvated the five complexes with TIP3P and a water buffer size of 12 Å and added 6 Cl^-^ ions to neutralize the formal charges of the system. After the standard setup detailed in Supporting Information (SI), production runs of the five systems were performed at 298K for 500 ns in NPT ensemble and saved every 2 ps with a 2-fs time step by using pmemd.cuda from the AMBER 14 package. For each CDK8/CycC–PL complex, we chose the equilibrated conformations at 298K and conformations after 100-, 400-, 500-ns MD simulations as initial structures to run 12 metadynamics simulations. Details of the collective variables chosen for metadynamics are in SI. All the trajectories and input files are available upon request.

### Computing free energy and unbinding time by using milestoning theory

To construct unbinding free energy and compute the kinetics properties with milestoning theory, we needed to define milestones along reaction coordinates and estimate the probability of a transition between two milestones by using many short classical MD simulations. The PCA obtained from a metadynamics trajectory of a complex was used to define the milestones. To obtain many short classical MD runs for each complex, we resaved a frame every 50 ps from a metadynamics and PSIM trajectory. As a result, about 500 to 600 conformations were used as initial conformations, and we ran 20 replicas of 100-ps classical MD simulations for each initial conformation with a 2-fs time step at 298K. Frames were saved every 50 fs for each 100-ps MD trajectory, and the settings were the same as the MD simulation mentioned previously.

To compute PCA for a metadynamics trajectory, we selected the α-carbon atoms of CDK8/CycC and heavy atoms of PLs, and computed the covariance matrix of the Cartesian coordinates of these atoms by using the first frame in each trajectory as references. We saved the eigenvectors of the covariance matrix and used the equation PC_i_ = R^T^ (X(t)-<X>) to project frames from the metadynamics trajectory onto PC1 and PC2 space, where R^T^ is the eigenvector with the highest eigenvalue for PC1 and second highest for PC2, and X(t) and <X> are the Cartesian coordinates of the selected atoms at time t and the average over the trajectory, respectively. Frames from a metadynamics trajectory were projected onto PC1 and PC2 space, as exemplified in Figure 8A. The original metadynamics trajectory was further smoothed by averaging forward and backward 100 frames, and the frames were also projected onto the PC space, as shown by the red line in Figure 8A. The smoothed trajectory can more clearly represent the dissociation path, and multiple short lines of 20.0 eigenvalue units in length were placed perpendicular to the path. Each line is about 3.0 eigenvalue units apart, and the lines were optimized to minimize the overlapping of the lines, as illustrated in Figure 8B. Therefore, each line serves as one milestone. The structures nearby the milestones were used to initiate short MD simulations that generated trajectories between neighboring milestones. Frames saved from the each 100-ps short MD run were then projected onto the PC1/PC2 space and fall into a space between neighboring milestones (Figure S12).

The average lifetime and transition time computed from the conformations that significantly deviate from the define pathway may be less meaningful, compared with those that computed from conformations close to the defined pathway. Based on this same reason, we removed the data points that fall onto the areas that were 10 units away from the path in PC1/PC2 space (Figure S12). The remaining data points were used to compute the duration of each milestone and the transition counts between adjacent milestones.

Finally, the transition kernel (matrix) *K*, free energy profile and residence time were computed following the milestoning theory. ^30, 32, 53^ In brief, the probability of a transition between two milestones *i* and *j* is *K*_*ij*_, the average lifetime of a milestone *i* is *t*_*i*_, and the number of trajectories that pass through milestone *i* in unit time is *q*_*i*_, which is also termed the stationary flux. The steady state *q*_*i*_ is given by a solution of the linear equations ∑_*i*_ *q*_*i*_ *K*_*ij*_ = *q*_*i*_. The stationary probability of milestone *i* can be approximated by *q*_*i*_*t*_*i*_ and then relates to the free energy *Gi* of trajectories that passed milestone *i, Gi* = −k_B_Tln(*q*_*i*_*t*_*i*_). The residence time of a ligand before passing milestone *f* is determined by *τ*_*f*_ = **P**^***t***^**·**(**I**−**K**)^***t***^**t**.

### Compound synthesis and kinetics assay

PL4-OH is synthesized by ChemConsulting LLC, and information of the synthesized compound is detailed in SI. Kinetic-assay testing was conducted by Proteros biostructures GmbH.^54^ In brief, the company uses the Proteros Reporter Displacement Assay which is based on reporter probes designed to bind to the binding site of CDK8/CycC, and the proximity between a reporter probe and protein results in the emission of an optical signal. When a test ligand binds to the target protein, the reporter probe is displaced, which leads to a signal. Monitoring the time dependent signal yields the binding kinetics of the ligand.

## Acknowledgments

This study was supported by the US National Institutes of Health (GM-109045), US National Science Foundation (MCB-1350401), and NSF national supercomputer centers (TG-CHE130009). We thank Dr. Ron Elber for discussion relating to milestoning theory and Dr. Ming Lee Tang and Tony Dorado for compound synthesis.

## Abbreviations

SRK: structure-kinetic relation
PCA: principal component analysis
H-bond: hydrogen bond
MD: molecular dynamics
PL: pyrazolourea ligand
CDK8: cyclin-dependent kinase 8
CycC: cyclin C
PSIM: pathway search guided by the internal motions

